# Antinociceptive, hypothermic, and appetitive effects of vaped and injected Δ9-tetrahydrocannabinol (THC) in rats: exposure and dose-effect comparisons by strain and sex

**DOI:** 10.1101/2020.10.06.327312

**Authors:** Catherine F. Moore, Catherine M. Davis, Eric L. Harvey, Michael A. Taffe, Elise M. Weerts

**Affiliations:** Division of Behavioral Biology, Department of Psychiatry and Behavioral Sciences, Johns Hopkins University School of Medicine, Baltimore, MD; Department of Psychiatry, University of California San Diego, La Jolla, CA; Department of Pharmacology and Molecular Therapeutics, Uniformed Services University of the Health Sciences, Bethesda, MD

**Author notes:** Corresponding author at: Johns Hopkins Bayview Research Campus, Behavioral Biology Research Center, 5510 Nathan Shock Drive, Suite 3000, Baltimore, MD 21224, USA. E-mail address (C.F. Moore).

**Keywords:** THC, Vapor Exposure, Cannabinoids, Nociception, Hypothermia, Appetite

## Abstract

Advances in drug vapor exposure systems utilizing e-cigarette technology have enabled evaluation of Δ-9-tetrahydrocannabinol (THC) vapor effects in laboratory animals. The purpose of this study was to 1) establish a range of parameters of THC vapor exposure in rats sufficient to produce a behavioral dose-effect curve in a battery of tasks sensitive to THC; 2) to investigate sex differences in the effects of THC vapor exposure and THC injection (intraperitoneal, IP) on these behaviors in two strains of outbred rats. Male and female Wistar and Sprague Dawley rats (N=22, 5-6/group) received THC via IP injection (1-20 mg/kg) and passive exposure to THC vapor (200 mg/ml; 5 conditions) in a within subject design. The effects of vaped and injected THC were determined using the tail-withdrawal assay for nociception, rectal measurements of body temperature, and progressive-ratio responding for food pellets. Plasma THC concentrations were assessed after 10 mg/kg IP THC or THC vapor. THC produced dose and exposure-dependent antinociception and hypothermia. THC vapor produced inverted U-shaped effects in motivation to obtain food, while IP THC reduced PR breakpoints. Plasma THC concentrations were higher after 10 mg/kg IP THC (152 ng/mL) compared to the highest vapor exposure condition tested (38 ng/mL). THC vapor exposure produces reliable, dose-orderly effects on nociception, body temperature, and food-maintained behavior that is comparable to effects observed after IP THC. There are considerable differences between the time course of behavioral outcomes produced by these two different routes of administration.

## 1. Introduction

Cannabis is one of the most widely used drugs in the world, including in the United States. It is most often used by inhalation, and more recently, vaping of cannabis and cannabis extracts containing ∆-9-tetrahydrocannabinol (THC, the primary psychoactive constituent of cannabis) is on the rise (Giroud et al., 2015; Ramo et al., 2015; Budney et al., 2015; Morean et al., 2015; Jones et al., 2016; Lee et al., 2016; Varlet et al., 2016; Trivers et al., 2019). Rates of cannabis vaping are estimated to be between 20%-37% (past 30-days) and 60% (lifetime) for cannabis users (Lee et al., 2016; Schauer et al., 2020; Goodman et al., 2020).

Although most preclinical studies of cannabis and cannabinoids have employed injection methods (intraperitoneal, subcutaneous, intravenous), preclinical vapor exposure models have more recently been developed using methods for aerosolizing THC (Lichtman et al., 2000; Wilson et al., 2002), use of desktop vaporizers (i.e. Volcano© vaporizer)(Manwell et al., 2014a), and e-vape technology (Nguyen et al., 2016); (for review, see Moore et al., 2020; Miliano et al., 2020). Route of administration is important from a translational perspective and for interpretation of pharmacokinetic and pharmacodynamic effects of cannabinoids.

The main objective of this study was to develop a range of vapor exposure conditions to functionally ‘anchor’ a behavioral dose-effect curve (i.e., range from no effect to response inhibition) and compare behavioral and physiological effects of THC vapor to intraperitoneally (IP) injected THC. In addition, the proposed studies were intended to establish the vapor exposure model in our laboratory at Johns Hopkins University and compare with similar studies examining sex and strain differences conducted elsewhere. THC vapor exposure parameters including puff duration (2-9s), puff frequency (5-20x), and total exposure time (10-30m) were varied to generate a range of vaporized e-liquid volumes (200 mg/ml THC). We administered a range of THC doses (1-20 mg/kg, IP) to generate a full dose-response curve for comparison with THC vapor in a behavioral test battery. Behavioral measures included a tail-withdrawal assay for nociception and rectal measurements of body temperature, and food-maintained operant responding. These behaviors were chosen based on the established effects of THC on antinociception, hypothermia, and appetite (Compton et al., 1993; Higgs et al., 2005; Metna-Laurent et al., 2017). Blood was collected and plasma THC concentrations were analyzed following 1 IP dose and 1 vapor exposure condition. We used a within-subject design in male and female Sprague-Dawley and Wistar rats to assess potential sex and strain differences in THC effects.

## 2. Methods

### 2.1 Subjects

Experiments were performed using adult male and female Wistar rats (N=12, 6 of each sex) and Sprague-Dawley rats (N=12, 6 of each sex) (Charles River, Wilmington, MA). Rats were single-housed in wire-topped, plastic cages (27 × 48 × 20 cm) on a 12-hour reverse light cycle (lights off at 9:00 a.m.), in an AAALAC-approved humidity-controlled and temperature-controlled vivarium. Male and female rats weighed 250-350 grams at the start of experiments and were maintained at 90% of their free feeding weight throughout the course of the experiments. Diet was a corn-based chow (Harlan Teklad Diet 2018; Harlan, Indianapolis, IN) and rats had free access to water at all times with the exception of during experimental test procedures. Procedures used in this study were approved by the Johns Hopkins Institutional Animal Care and Use Committee and adhered to the National Institutes of Health *Guide for the Care and Use of Laboratory Animals*. Two rats were euthanized during the course of experiments for health reasons unrelated to the treatments; therefore, final group sizes were Sprague Dawley (N=6 males, N=5 females) and Wistar (N=6 males, N=5 females).

### 2.2 Drugs

THC stock solution (200 mg/ml in 95% ethanol) was provided by the U.S. National Institute on Drug Abuse Drug Supply Program. For vapor administration, the 200 mg/ml ethanol-based THC stock solution was mixed in 100% propylene glycol and the ethanol was evaporated using nitrogen to yield a 200 mg/ml THC solution for vaporization. For intraperitoneal (IP) injections, the THC stock was dissolved in a vehicle solution of 0.9% sterile saline, ethanol, and Cremophor EL (18:1:1 ratio) for final doses of 1, 3, 5.6, 10, and 20 mg/ml, administered at 1 ml/kg. Initial test doses were selected based on the literature (Taffe et al., 2015; Craft et al., 2019).

### 2.3 Vapor exposure system

A commercial vapor chamber system (La Jolla Alcohol Research Institute, La Jolla CA) was utilized. The system contained four sealed polycarbonate rat cages (35 × 28 × 26 cm; 25L) adapted for the delivery of vaporized drug, an electronic vapor device (e-vape) with connecting tubes and an air pump to regulate continuous airflow. The chamber air was vacuum controlled by the air pump which pulls room ambient air into the chamber through an intake valve and out through the exhaust valve at a constant rate of 2-3 L/minute. The e-vape device (2^nd^ generation) was a Smok Baby Beast Brother TFV8 Sub-Ohm Tank (with the V8 X-Baby M2 0.25-Ω coil; SMOKTech, Nanshan, Shenzhen, China). The e-vape controller was set to a maximum temperature of 400°F (~30W).

### 2.4 Vapor Delivery System Evaluation

Vapor delivery systems were tested to determine volumes delivered under different puff durations and frequencies. The amount of e-liquid (mL) vaporized per vapor delivery and condition was determined by weighing of the vapor tank before and after vapor deliveries of the propylene glycol vehicle over repeated testing. Each condition was tested in triplicate in a randomized order.

### 2.5 Study design

A randomized, within subject design was utilized for the study. Test sessions were conducted weekdays (Mon-Fri), with one vehicle test and one drug test conducted each week, and a minimum of at least 7 days between each THC dose for each individual subject. Weekly vehicle tests were included to control for any shifts in baseline behaviors over time. Each week, half of the rats received vapor and half received IP injections, and the route of THC administration alternated weekly until full dose-response curves were generated for each subject for both vapor and IP injections. THC injections were administered in a blind, within-subject Latin-square design (0-20 mg/kg). For the vapor exposure testing, the number of puffs, duration of puffs, and inter-puff intervals were systematically increased to produce 5 vapor conditions as detailed in Table 1, tested in ascending order. Vapor Condition 1 was repeated in the middle of the testing period (week 8) to assess any changes in response to THC vapor exposure. The total testing period was approximately 14 weeks.

**Table 1.**
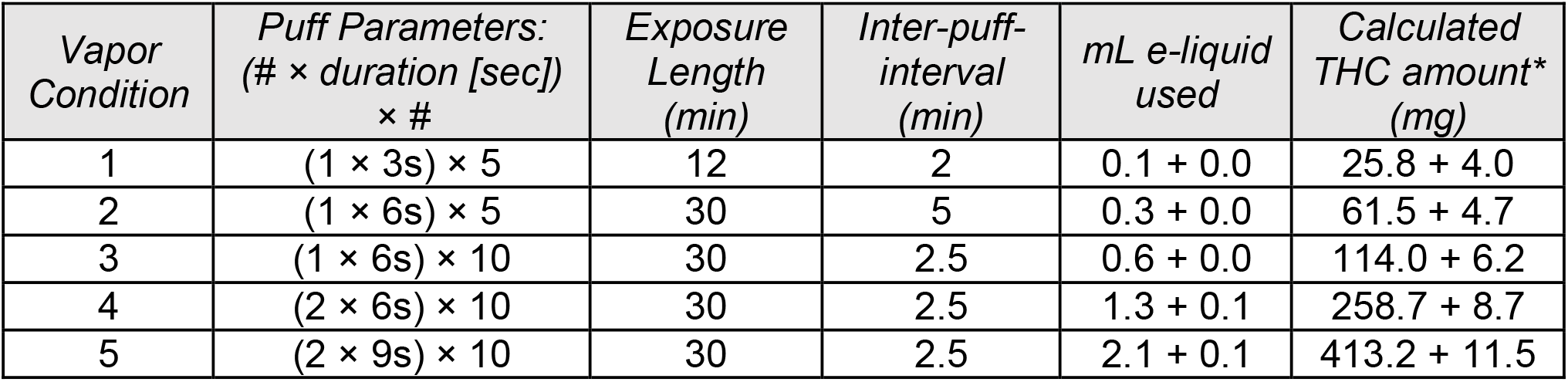
Summary of parameters used in vapor exposure conditions. For conditions 4 and 5 a series of 2 puffs was delivered in rapid succession with 2s in between to mitigate overheating of the vapor coil. Following the last vapor delivery of each condition, rats remained in the cage to allow enough time (e.g., 4-10 min) for vapor exposure and clearance of vapor from the chamber prior to testing. *Based on 200 mg/ml concentration used in these studies; this amount of vaporized drug is divided evenly between 4 chambers.

| *Vapor Condition* | *Puff Parameters:*<br>*(# × duration [sec])*<br>*× #* | *Exposure Length*<br>*(min)* | *Inter-puff-interval*<br>*(min)* | *mL e-liquid used* | *Calculated THC amount\**<br>*(mg)* |
| --- | --- | --- | --- | --- | --- |
| 1 | (1 × 3s) × 5 | 12 | 2 | 0.1 + 0.0 | 25.8 + 4.0 |
| 2 | (1 × 6s) × 5 | 30 | 5 | 0.3 + 0.0 | 61.5 + 4.7 |
| 3 | (1 × 6s) × 10 | 30 | 2.5 | 0.6 + 0.0 | 114.0 + 6.2 |
| 4 | (2 × 6s) × 10 | 30 | 2.5 | 1.3 + 0.1 | 258.7 + 8.7 |
| 5 | (2 × 9s) × 10 | 30 | 2.5 | 2.1 + 0.1 | 413.2 + 11.5 |

### 2.6 Behavioral Test Battery

Subjects completed a battery of behavioral tests with repeated measures over a 5-hour test period. The sequence and time course of testing are depicted in **Figure 1**. Below, we detail methods for each procedure in the behavioral test battery.

**Figure 1.**
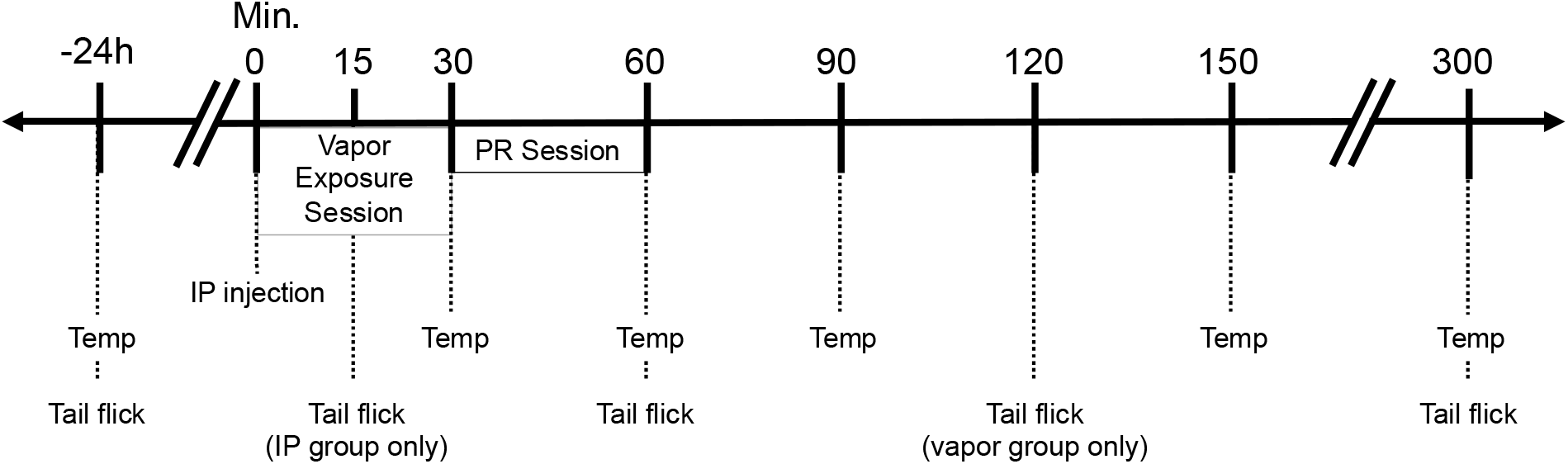
Schematic of drug or vehicle administration and time course of measurements of rectal temperature, tail withdrawal latency and food-maintained responding in Progressive Ratio (PR) Sessions in the behavioral test battery. Time noted is from injection or vapor initiation. Note: Vapor Condition 1 lasted only 12 minutes and subsequent testing took place on the same schedule, therefore the true time-points for this condition only are off by 18-min (e.g. 30-min time point is actually only 12-min from vapor initiation and so on).

#### 2.6.1 Antinociception (thermal pain sensitivity)

Thermal pain sensitivity was assessed using the tail withdrawal (TW) assay. In this test, the distal end of the rat’s tail (~50 mm from the tip) is exposed to radiant heat from a precise photobeam (Harvard Apparatus, Cambridge, MA, USA). Latency to respond to the heat stimulus by flexion of the tail is recorded. Prior to testing the radiant heat setting was calibrated to achieve a baseline latency of 4s. Baseline TW latencies were obtained 24-hrs prior to drug administration. Rats were then tested at 3 time points post-drug administration on the test day, 15-min (IP only), 60-min, 120 min (vapor only), and 300 min.

#### 2.6.2 Body temperature

Body temperature was determined with a digital rectal thermometer with a lubricated flexible probe. Baseline body temperature was obtained 24-hrs prior to drug administration (i.e. same time of day as injection/vapor initiation). Changes from baseline temperature were evaluated in comparison to vehicle controls across 5 time points on the test day (30, 60, 90, 150, 300 min).

#### 2.6.3 Food-maintained Operant Responding

##### Training

Thirty minute operant test sessions occurred daily (Mon-Fri) in dedicated experimental chambers interfaced with a personal computer with Med-PC hardware and software for experimental control of behavior (Med Associates, St. Albans, VT). Each experimental chamber was equipped with a nose-poke key, a cue light, and a food cup connected to an automated pellet feeder for delivery of food pellets, all positioned inside a sound-attenuating enclosure with exhaust fans. Rats were initially trained to press a nose poke key for delivery of a 45-mg food pellet under a fixed ratio 1 (FR1) schedule of reinforcement. After stability under an FR1 schedule (responses ±10%), the FR requirement was gradually increased 10 (FR10).

##### Progressive ratio

After stability of responding was obtained under the FR10, rats were shifted to a progressive ratio (PR) procedure, in which the number of responses required to produce a food pellet (ratio requirement) was progressively increased (1, 2, 4, 6, 9, 12, 15, etc.)(Richardson and Roberts, 1996). The last completed ratio requirement resulting in reinforcement before the animal stops responding within a specified duration (10-min), or before the maximum session time of 30 minutes was reached, was defined as the 'break point' (Richardson and Roberts, 1996). PR break points for food pellets were assessed immediately after the end of THC exposure and 30 minutes after IP administration. Response rates during the session were calculated as total responses/total responding time and used as an indicator of motor impairment. Non-drug PR sessions were continued on subsequent days to ensure that rats were maintaining stable responding for food.

### 2.7 Blood Collection Procedures

At the end of the study, selected conditions (vapor condition 5; 10 mg/kg, IP) were repeated for blood sampling. Blood samples (~250 μl) were collected from the saphenous vein in unanesthetized rats, 30 minutes after IP injection or immediately following the end of the vapor exposure session. Blood was collected into EDTA-coated microcentrifuge tubes, placed on wet ice for 30 minutes, and then centrifuged at 3000G for 10 minutes. Plasma supernatant was transferred to low protein binding microcentrifuge tubes and stored at −80°C until analysis.

### 2.8 Plasma THC analysis

Plasma THC concentrations were quantified using fast liquid chromatography/mass spectrometry (LC/MS) adapted from (Lacroix and Saussereau, 2012; Irimia et al., 2015; Nguyen et al., 2018). 50 μL of plasma were mixed with 50 μL of deuterated internal standard (100 ng/mL CBD-d3 and THC-d3; Cerilliant), and cannabinoids were extracted into 300 μL acetonitrile and 800 μL of chloroform and then dried. Samples were reconstituted in 100 μL of a methanol/water (2:1) mixture. Separation was performed on an Agilent LC1100 using an Poroshell 120 EC-C18 column (4.0μm, 2.1mm × 100mm) using isocratic elution with water and methanol, both with 0.2 % formic acid (250 μL/min; 81% MeOH). THC was quantified using an Agilent 6140 single quadrupole MSD using electrospray ionization and selected ion monitoring [THC (m/z=315.2) and THC-d3 (m/z=318.2)]. Calibration curves were conducted daily for each assay at a concentration range of 0-200 ng/mL and observed correlation coefficients were 0.999.

### 2.9 Data Analysis

Outcome measures submitted for analysis included: body temperature (°C), TW latency percent of maximum possible effect [%MPE], PR break points, PR response rate, and plasma THC concentration (ng/mL). Antinociception was calculated as percent of maximum possible effect (% MPE= [(test latency– baseline latency)/(maximum latency – baseline latency)] × 100). Baseline body temperature and TW latencies were obtained 24-hr before drug testing days. Breakpoints were determined by the response requirement in effect when an animal stopped responding. Response rate under the PR schedule of reinforcement was calculated as total responses/total responding time.

For each route of administration (vapor exposure and IP administered), three or four-way repeated measures ANOVAs were conducted with strain and sex as between subjects variables and vapor condition/dose and time after the start of inhalation or after injection (when applicable) as within subject variables. Next, two or three-way ANOVAs were conducted within each strain, with sex as a between subjects variable and vapor condition/dose and time (when applicable) as within subject variables. To evaluate main effects or interactions, post-hoc one- or two-way ANOVAs were conducted with strain as a between subjects effect (collapsed across sex), and vapor condition/dose and time (when applicable) as within-subjects effects; as well as within each sex (strains were never collapsed) with vapor condition/dose and time (when applicable) as within-subjects effects. Dunnett’s post-hoc tests were used to analyze differences in outcomes between dose/condition and vehicle. If a main effect of or interaction with strain or sex was indicated, sex or strain differences were determined with a one or two-way ANOVA within each dose (effects of Sex and Time, when applicable) and Sidak’s post-hoc tests were used. Two-sided unpaired t-tests were used as post-hoc tests to evaluate sex differences in plasma THC levels. Statistics were performed in Statistica 11 (Stat Soft, Inc.) and GraphPad Prism. The alpha level was set at 0.05 for significance. All ANOVA results are reported in Supplementary Tables 1-5 and the highest order effect/interaction is reported in the text.

## 3. Results

### 3.1 Vapor Delivery System Evaluation

The vapor delivery system reliably and systematically produced vapor from the e-liquid (mL) with the amount used related to the puff duration (**Figure 2A**). The amount of e-liquid vaporized by the conditions used in the study ranged from 0.1-2.1 mL (25-413 mg THC; 200 mg/mL concentration; see Table 1 and **Figure 2B**).

**Figure 2.**
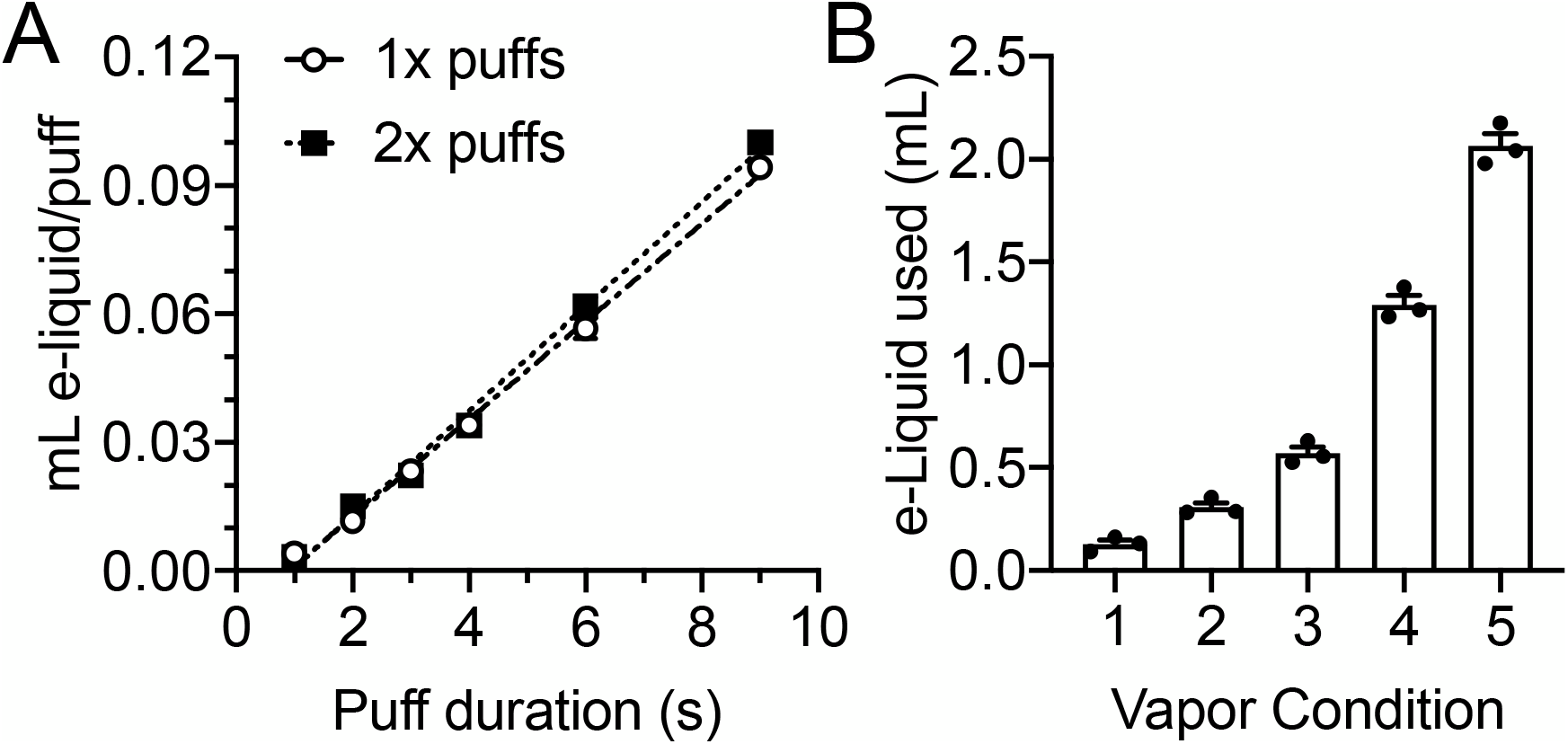
Amount of e-liquid vaporized per puff in a series of 1x or 2x puffs of different durations (A) and total e-liquid used per each vapor exposure condition (1-5)(B). Data are Mean ± SEM.

### 3.2 THC effects on TW latency

THC vapor exposure increased TW latency, a measure of thermal nociception (**Figure 3A**), as confirmed by a significant interaction of Vapor Condition × Time (F(10, 180) =6.43, p<0.001). In Sprague-Dawley rats, there were significant main effects of Vapor Condition (F(5, 45) =20.37, p<0.001) and Time (F(2, 18) =3.64, p<0.05) on TW latency. In the Wistar rats there were significant interactions between Time × Sex (F(2, 18) =3.94, p<0.05) and Vapor Condition × Time (F(10, 90) =6.24, p<0.001) on TW latency. In both Sprague-Dawley and Wistar rats, only the highest THC vapor conditions (4-5) increased antinociception during one or more time points when compared to vehicle vapor. Peak antinociceptive effects occurred at the earliest time point tested (60 min for Vapor Conditions 4 and 5, see Supplemental Table 6).

**Figure 3.**
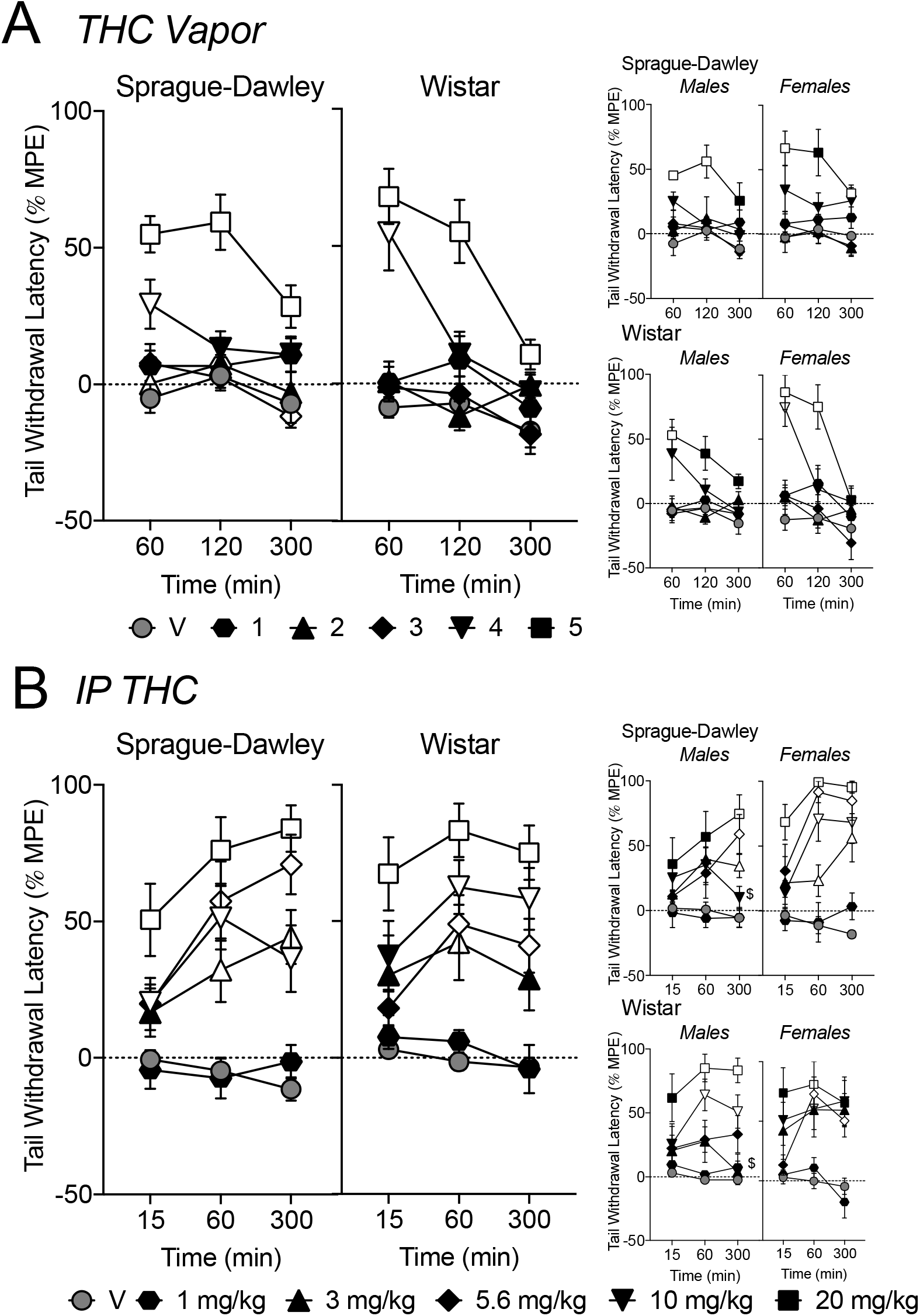
[Top panel] Tail withdrawal latencies, expressed as % of maximum possible effect (%MPE) after exposure to THC (200 mg/ml; conditions 1-5) or the propylene glycol vehicle (V) vapor. Males and females are shown separately in the right panel. [Bottom panel] Tail withdrawal latencies (%MPE) after IP THC (1-20 mg/kg) or vehicle (V). Males and females are shown separately in the right panel. Data are Mean ± SEM. Symbol colors: Vehicle symbols are dark gray; other symbols are drug, and unfilled symbols denote a difference from Vehicle (p<0.05).

IP THC increased TW latency (**Figure 3B**), evident in a significant 3-way interaction of Dose × Time × Sex (F(10, 180) =2.08, p<0.05) on TW latency. In Sprague-Dawley rats, there was a significant interaction of Dose × Time × Sex (F(10, 90) =2.29, p<0.05) on TW latency. Female Sprague-Dawley rats showed antinociceptive effects at lower doses of THC, and these effects appeared earlier after injection than in male rats, with significant effects occurring from 60-300 min (3-20 mg/kg), while the male Sprague-Dawley rats did not show significant increases in TW latency until 300 min. In Wistar rats, there were significant main effect of Dose (F(5, 45) =21.77, p<0.001) and Time (F(2, 18) =6.57, p<0.01) on TW latency. In both Sprague-Dawley and Wistar rats, almost all THC doses tested (3-20 mg/kg), with the exception of the lowest dose, increased TW latency during one or more time points when compared to vehicle vapor. Peak antinociceptive effects after most THC doses occurred at later time points (60-300 min), while the highest THC dose caused peak antinociceptive effects earlier (between 15-60 min; see Supplemental Table 7).

### 3.3 THC effects on body temperature

THC vapor exposure decreased body temperature (**Figure 4A**), confirmed by significant 3-way interactions of Time × Strain × Sex (F(4, 72) =4.20, p<0.01), Vapor Condition × Time × Strain (F(20, 360) =2.76, p<0.001), and Vapor Condition × Time × Sex (F(20, 360) =1.66, p<0.05) on body temperature. In Sprague-Dawley rats, there were significant interactions between Time × Sex (F(4, 36) =2.67, p<0.05) and Vapor Condition × Time (F(20, 180) =6.30, p<0.001) on body temperature. In Wistar rats, there was a significant 3-way interaction between Vapor Condition × Time × Sex (F(16, 144) =2.09, p<0.05) on body temperature. In Sprague-Dawley rats, all THC vapor conditions (1-5) reduced body temperature during one or more time points when compared to vehicle vapor. At 300 minutes, body temperatures had mostly returned to control levels except for vapor conditions 2 and 5. In Wistar rats, THC vapor conditions 3-5, but not 1-2, reduced body temperature during one or more time points when compared to vehicle vapor, all returning to control levels by 300 minutes. Maximal hypothermic effects occurred after the highest THC vapor condition (−0.54°C in Sprague Dawley and −0.70°C after in Wistar rats, see Supplementary Table 6). The temperature nadir occurred 60 min post vapor-initiation.

**Figure 4.**
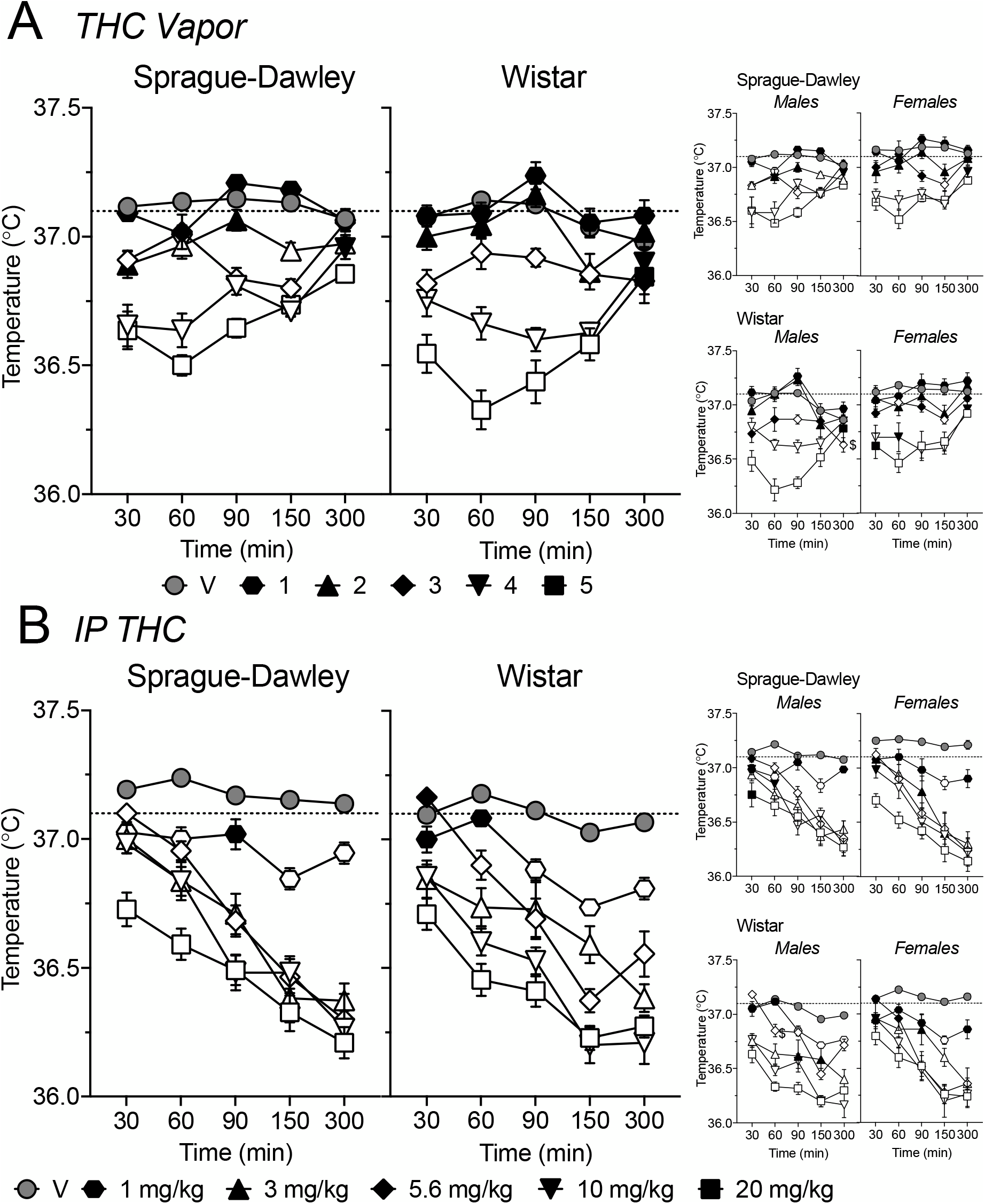
[Top panel] Body temperature (°C) after exposure to THC (200 mg/ml; conditions 1-5) or the propylene glycol vehicle (V) vapor. Males and females are shown separately in the right panel. [Bottom panel] Body temperature (°C) after IP THC (1-20 mg/kg) or vehicle (V). Males and females are shown separately in the right panel. Reference line reflects the overall average temperature under vehicle conditions (37.1°C). Data are Mean ± SEM. Symbol colors: Dark gray symbols are Vehicle; other symbols are drug, and unfilled symbols denote a difference from Vehicle (p<0.05).

IP THC also decreased body temperature (**Figure 4B**) as confirmed by a significant three-way interaction of Dose × Time × Strain (F(20, 360) =2.86, p<0.001) on body temperature. In Sprague-Dawley rats, there were significant interactions between Dose × Sex (F(5, 45) =2.78, p<0.001) and Dose × Time (F(20, 180) =7.89, p<0.001) on body temperature. In Wistar rats, there was a significant 3-way interaction between Dose × Time × Sex (F(20, 180) =2.39, p<0.05) on body temperature. In both Sprague-Dawley and Wistar rats, all THC doses (1-20 mg/kg) reduced body temperature during one or more time points when compared to vehicle. In Wistars, lower doses (1 and 5.6 mg/kg) took longer to reduce body temperature (decreases at 90 and 60 minutes, respectively). Wistars showed larger hypothermic effects to 10 mg/kg THC (significant differences from Sprague-Dawley rats at 60 and 150 min). Maximal hypothermic effects occurred after the two highest THC doses, 10 and 20 mg/kg (−0.99°C in Sprague Dawley after 20 mg/kg THC and −0.98°C after 10 mg/kg in Wistar rats). Hypothermic effects of IP injected THC continued to increase over the testing period, with the temperature nadir observed at the last time-point tested (300 min).

### 3.4 THC effects on PR break points for food

THC vapor exposure modulated PR break points in a condition and strain-dependent manner (**Figure 5A**), confirmed by a significant interaction between Vapor Condition × Strain (F (5, 90) = 2.53, p<0.05) on PR break points. Particularly, in Sprague-Dawley rats, THC vapor produced biphasic effects on PR break points (Vapor Condition: F (5, 45) = 20.20; p<0.001). PR break points were increased after the lower vapor exposure conditions (1 and 2) and decreased after the two highest vapor exposure conditions (4 and 5; p’s<0.05). While ANOVA did not indicate any main effect of (or interactions with) sex, the increased PR break points seen after vapor condition 1 appear to be driven by female rats (p<0.05), while male rats show a trend for increased PR breakpoints after vapor condition 2 (p=0.07). In Wistar rats, THC vapor exposure reduced PR break points (Vapor Condition: F (5, 45) = 20.57; p<0.001). PR break points were decreased after the two highest vapor exposure conditions (4 and 5; p’s<0.05). THC vapor condition 1 was repeated in the middle of the testing period (week 8) to assess any changes in effects after repeated exposure to THC. PR breakpoints following THC vapor condition 1 were equivalent when tested in week 1 and week 8 (no effect of test week: F (1, 19) = 2.476, *n.s.*). An average of these two weeks was used in the above described analysis.

**Figure 5.**
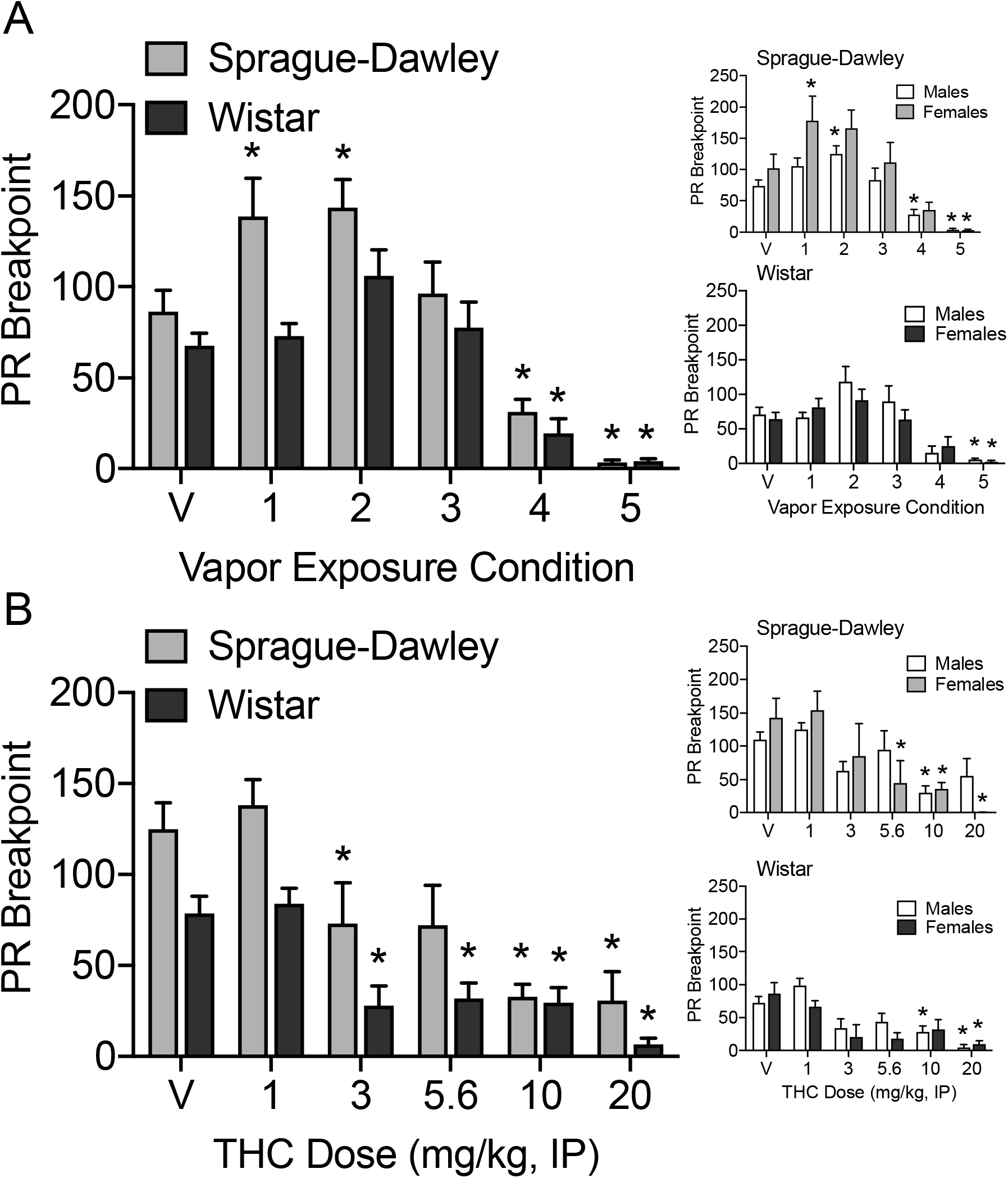
[Top panel] PR breakpoints after exposure to THC (200 mg/ml; conditions 1-5) or the propylene glycol vehicle (V) vapor. [Bottom panel] Mean PR breakpoints after IP THC (1-20 mg/kg) or vehicle (V). Males and females are shown separately in the right panel. Data are Mean ± SEM. Asterisks (*) denote difference from Vehicle (p<0.05) within the respective group.

IP THC reduced PR breakpoints in all rats (**Figure 5B**). Overall, there were significant main effects of Dose (F(5, 90) = 17.69, p<.0001) and Strain (F(1, 18) = 16.34, p<0.001) on PR break points. In Sprague-Dawley and Wistar rats, there was a significant main effect of Dose (Sprague-Dawley: F(5, 45) = 8.39, p<0.001; Wistar: F(5, 45) = 13.20, p<0.001) and in Sprague-Dawley rats, there was also a significant main effect of Sex (F(1, 45) = 6.63, p<0.05), though no post-hoc tests comparing males and females at each dose were significant. In both Sprague-Dawley and Wistar rats, PR break points were reduced after 3-20 mg/kg THC.

PR breakpoints assessed each week after vehicle administration (IP and vapor exposure) showed no change over the course of the testing period. (Vehicle Vapor, no effect of test week: F(4, 80) = 0.29, *n.s.*); Vehicle IP, no effect of test week F(6, 120) = 1.04, n.s.).

There was an effect of THC (vapor and IP) on response rate under the PR schedule of reinforcement (Vapor Condition: F (5,100) = 30.08, p<0.0001; Dose: F (5,100) = 7.58, p<0.0001; data not shown). There was also a main effect of sex on IP THC effects on response rate (Sex (F(1,18) =4.81, p<0.05), but no effects of sex on THC vapor effects, and no main effects of strain, on response rate. In Sprague-Dawley rats, vapor conditions 3-5 decreased response rate compared to vehicle vapor, and 10-20 mg/kg THC decreased response rate compared to IP vehicle. In Wistar rats, vapor conditions 4-5 decreased response rate compared to vehicle vapor, but post-hoc tests did not indicate significant differences from IP vehicle between response rates after any IP THC doses. These reductions in response rates after high doses of THC are indicative of a motor suppressive, rather than a motivational or appetitive effect.

### 3.5 Plasma THC following injection and vapor exposure

One plasma sample was determined to be an outlier (>15 standard deviations from jackknifed group mean) and removed (1 Sprague-Dawley female, 10 mg/kg IP). Following THC vapor exposure, there were no effects of strain or sex on plasma THC concentrations (**Figure 6A**). Following injection of 10 mg/kg THC, there was an effect of Sex (F(1, 17)=5.34, p<0.05) but not strain on plasma THC concentrations, with Sprague-Dawley females showing higher plasma THC compared to males of the same strain (**Figure 6B**). Plasma THC concentrations following 10 mg/kg IP THC (30-min after injection) were 4-fold higher than observed after THC vapor exposure (10 minutes following removal from vapor chamber).

**Figure 6.**
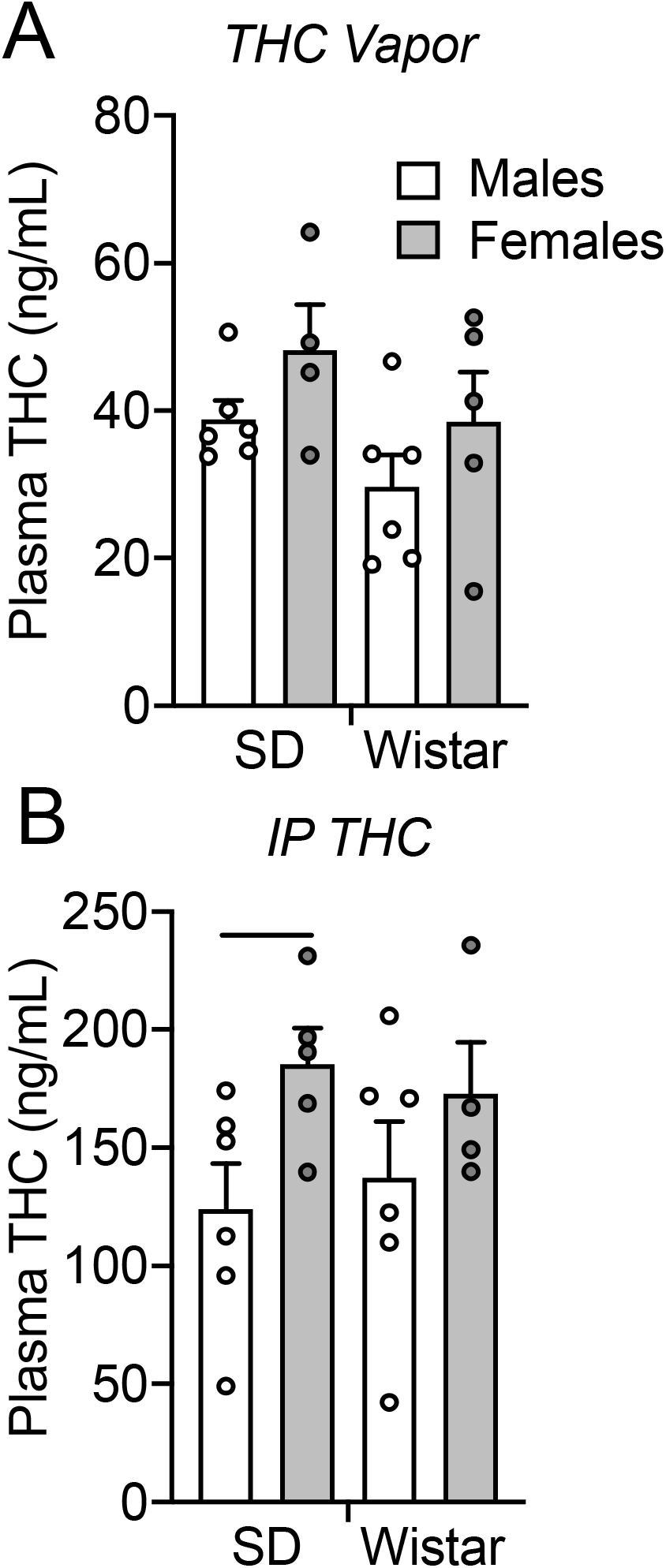
Plasma THC concentrations (ng/mL) following THC vapor exposure in Condition 5, 20 x 9s puffs of 200 mg/ml THC (A) or 10 mg/kg IP THC. Blood collection occurred 30-min from injection or immediately (approx. 10-min) after a 30-min THC vapor exposure session. Data are Mean ± SEM. Asterisks (*) denote sex difference (p<0.05).

## 4. Discussion

This series of experiments aimed to validate a passive THC vapor exposure model in rats and compare exposure/dose-response relationships in two different routes of administration across sex and rat strain. Our vapor exposure model produced consistent, reliable and puff-orderly amounts of vapor and substantial plasma THC concentrations were detected after THC vapor exposure. We observed behavioral (nociception, appetite) and physiological (temperature) effects of THC vapor exposure (200 mg/ml; conditions 1-5) and IP THC (1-20 mg/kg) that differed with respect to route of administration. Below, we discuss each of these findings.

In the present study, we observed orderly antinociceptive effects of THC in a dose (IP) and exposure (vapor) dependent manner. Following IP THC (3-20 mg/kg), we observed antinociceptive effects that continued to increase in magnitude, with peak antinociceptive effects occurring at the last time point tested (300-minutes after injection). We also observed antinociceptive effects of THC vapor at the two highest exposure conditions tested (4-5). Peak antinociceptive effects of THC vapor exposure were observed between 60 and 120-min after vapor exposure initiation, with effects dissipating by 300-min after exposure except in the highest exposure condition (condition 5). The antinociceptive effects of THC vapor observed in the present study are similar to those reported by other labs utilizing IP administration and vapor exposure models (Tseng and Craft, 2001; Nguyen et al., 2018; Craft et al., 2019). For example, a study using Wistar rats found antinociceptive effects after 30-min vapor exposure to 200 mg/ml THC (Nguyen et al., 2018).

The current study also evaluated potential strain and sex differences. In Sprague-Dawley rats, after IP THC we observed an interaction between sex, time, and dose on antinociceptive effects in the tail withdrawal assay. Specifically, greater peak antinociceptive effects of THC were observed in Sprague-Dawley females compared to males at lower doses and the antinociceptive effects of IP THC occurred earlier in females. These data are consistent with studies by Craft and colleagues, in which greater antinociceptive effects of IP THC were observed in Sprague-Dawley females in a tail withdrawal assay and a paw pressure test (Tseng and Craft, 2001; Craft et al., 2019). Interestingly, in the present study, there were no sex differences in the antinociceptive effects of vaped THC in Sprague-Dawley rats. In Wistar rats, the opposite was observed, where no sex differences were seen after IP THC, but after vaped THC, female Wistar rats showed higher peak %MPE in the tail withdrawal (condition 5). Another THC vapor exposure study using Wistar rats found no difference in antinociceptive effects between males and females after 30-min exposure to 200 mg/ml THC (Nguyen et al., 2018), while in a study by Javadi-Paydar et al. (2018), Wistar males appeared to be less sensitive to THC.

THC also caused exposure/dose-dependent hypothermia after vapor exposure and IP injection. IP THC caused hypothermic effects that increased across the 5-hr testing period, resulting in roughly 1°C decreases after 20 mg/kg, observed 150-300 min post injection. The highest THC vapor condition produced maximal temperature decreases of 0.5-0.7°C that occurred at 60-min after vapor initiation, with body temperatures returning to control levels by 300-min. Multiple studies from the Taffe laboratory have reported hypothermic effects of IP THC and THC vapor exposure (Nguyen et al., 2016; Javadi-Paydar et al., 2018; Nguyen et al., 2020). In these studies, at the highest concentrations tested (100-200 mg/mL; 30-40 minutes exposure), temperatures decreased roughly 2.5 °C, which was comparable to temperature decreases (3°C) following 20 mg/kg, IP THC (Nguyen et al., 2016; Javadi-Paydar et al., 2018). Those prior studies report a similar time course of the hypothermic effects of THC vapor; in male and female Wistar rats, peak hypothermic effects occurred at 30-60 minutes after THC vapor initiation and were back to control levels after 4 hours (Javadi-Paydar et al., 2018; Nguyen et al., 2018; Nguyen et al., 2020). Also similar to the current study, they observed temperature nadir 4-5 hours after IP injection of THC (Nguyen et al., 2016), which is in stark contrast to the time course of hypothermic effects observed after THC vapor exposure.

The present study observed THC exposure/dose-dependent effects on appetite, as measured by PR break points. IP THC (3-20 mg/kg) produced dose-dependent decreases in PR break points. At the highest doses (10-20 mg/kg), response rates were also significantly reduced, indicating motor suppressive effects. We observed an inverted U-shaped effect of THC vapor exposure on PR break points for food, where low exposure conditions (1-2) increased PR break points, and high exposure conditions (4-5) decreased break points. Similar to IP THC, the highest vapor exposure conditions also caused reductions in response rates, indicating motor suppressive effects. Other studies using a Volcano© vaporizer for THC vapor exposure have observed stimulating effects on feeding and appetite: 10 mg THC increased food intake of a plain chow diet in the first hour of a 4-hr test (Manwell et al., 2014b) and vaporized cannabis plant material (7.8% (~62.4 mg) THC) increased free feeding of chow and palatable food in sated rats (Brutman et al., 2019). Inverted U-shaped dose effects of IP THC on PR break points for food have been reported in some studies and not others. For example, low doses of IP THC (1-3 mg/kg) increased breakpoints and higher doses (>5 mg/kg, IP) decreased break points for food (Solinas and Goldberg, 2005; Higgs et al., 2005). However, other studies show no effects (1-3 mg/kg) or dose-dependent decreases in PR breakpoints for food pellets following IP THC without observing increases (Olarte-Sanchez et al., 2015). We did not observe any increases of PR break points at low doses of IP administered THC. The discrepancy between these studies may be due in part to the macronutrient composition of the food pellet reinforcers. Chow pellets, such as those used in the present study as well as Olarte-Sanchez et al. (2015), are considered to be less palatable than food pellets higher in sucrose composition, such as those used in Solinas and Goldberg (2005). The ability of THC and other CB1 agonists to increase appetite is particularly evident in more palatable food (i.e. energy-dense, high in fat and/or sugar) (Koch, 2001; Berry and Mechoulam, 2002; Jager and Witkamp, 2014). In the present study, THC increased PR break points for plain chow in animals after vapor exposure only. As we did not test lower doses of IP THC (<1 mg/kg) we can’t rule out the possibility that a lower dose would not increase PR breakpoints. An analysis of sex differences in response to THC effects on PR breakpoints did not produce any significant main effects. However, disaggregating the data and analyzing the sexes separately, Sprague-Dawley female rats showed an increase in PR breakpoints after vapor condition 1, while Sprague-Dawley male rats showed an increase in PR breakpoints after vapor condition 2, suggesting females may be more sensitive to the appetite-enhancing effects of THC vapor.

Plasma THC following an injection of 10 mg/kg THC was, on average, 150 ng/ml 30-min after injection. This is consistent with other studies; for example 10 mg/kg IP THC produced plasma concentrations of 162 ng/ml THC in male rats (Nguyen et al., 2016). Our highest vapor exposure condition (5) produced ~38 ng/ml plasma THC immediately following removal from the vapor exposure session. Other THC vapor exposure studies report similar plasma THC concentrations; for example 30-min exposure to 100 mg/ml produced plasma THC concentrations of 67 ng/ml (Nguyen et al., 2019). Other studies using 200 mg/ml THC have observed higher plasma THC concentrations (e.g. 150-360 ng/ml) at this time point (Nguyen et al., 2018; Nguyen et al., 2020). However, multiple factors contribute to total exposure which have to be considered when making comparisons between laboratories, including exposure parameters and chamber set-up. For example, in the Taffe lab, where plasma THC concentrations have reached >300 ng/ml, the exposure chamber was much smaller than the ones employed in the present study (9 vs. 25L), the chamber design was 1 e-vape connected to 1 chamber (vs. 1 e-vape connected to 4 chambers, as in the current study), and air flow is discontinued in between puffs to allow for greater exposure (Nguyen et al., 2016; Nguyen et al., 2018; Nguyen et al., 2020). Interestingly, though plasma THC concentrations in the current study were considerably lower following vapor exposure compared to 10 mg/kg IP THC, behavioral outcomes were similar to the IP doses tested.

In summary, the present study advances vapor exposure methodologies, demonstrating a reliable and orderly exposure-effect curve with similarities and differences to the outcomes observed after IP THC. Importantly, there are considerable differences between the time course of behavioral outcomes produced by these two different routes of administration. Continued validation of vapor exposure methods and comparison of effects with those produced by routes of administration are important for approximate dose response curve comparisons. Furthermore, establishing parameters in vapor exposure needs to be done systematically to increase reproducibility between labs. Research on vapor methods are expanding and recently, vapor self-administration of cannabis extract was demonstrated in rats (Freels et al., 2020). Clearly, additional research using both passive exposure and response contingent vapor methods are important from a translational perspective and will further advance our understanding of the behavioral pharmacology vaped THC and other cannabinoids in cannabis.

## Acknowledgements

This work was supported by the National Institute on Drug Abuse of the National Institutes of Health grant numbers R21DA046154 (CD/EW) and R01DA042211 (MT), the Johns Hopkins University Dalio Fund in Decision Making and the Neuroscience of Motivated Behaviors (EW), and the Tobacco Related-Disease Research Program grant numbers TRDRP; T31IP1832 (MT). The authors also wish to thank Maury Cole and La Jolla Alcohol Research Inc. for development of custom vapor chamber systems and technical assistance. The opinions and assertions expressed herein are those of the author(s) and do not necessarily reflect the official policy or position of the Uniformed Services University or the Department of Defense.

