## Supplementary Tables for "Antinociceptive, hypothermic, and appetitive effects of vaped and injected Δ9-tetrahydrocannabinol (THC) in rats: exposure and dose-effect comparisons by strain and sex"

|  | TF Latency (%MPE) | | | | | | | | | | | |
| --- | --- | --- | --- | --- | --- | --- | --- | --- | --- | --- | --- | --- |
|  | (a) Vapor Exposure | | | | | | (b) IP injection | | | | | |
|  | all | | SD | | Wistar | | all | | SD | | Wistar | |
|  | F (df) | p | F (df) | p | F (df) | p | F (df) | p | F (df) | p | F (df) | p |
| **Strain** | 1.81  (1, 18) | 0.20 |  |  |  |  | 0.35 (1,18) | 0.56 |  |  |  |  |
| **Sex** | 2.06  (1, 18) | 0.17 | 0.91  (1, 9) | 0.37 | 1.17  (1, 9) | 0.31 | 3.97 (1,18) | 0.06 | 3.04  (1, 9) | 0.12 | 1.16  (1, 9) | 0.31 |
| Strain × Sex | 0.004  (1, 18) | 0.95 |  |  |  |  | 0.21 (1,18) | 0.65 |  |  |  |  |
| **Condition/Dose** | 39.66 (5, 90) | <0.001* | 20.37  (5, 45) | <0.001* | 19.82  (5, 45) | <0.001* | 45.83 (5,90) | <0.001* | 25.43  (5, 45) | <0.001* | 21.76  (5, 45) | <0.001* |
| Condition/Dose × Strain | 0.46  (5, 90) | 0.81 |  |  |  |  | 1.28 (5,90) | 0.28 |  |  |  |  |
| Condition/Dose × Sex | 1.85  (5, 90) | 0.11 | 1.18  (5, 45) | 0.34 | 1.03  (5, 45) | 0.41 | 2.33 (5,90) | 0.048* | 2.48  (5, 45) | 0.045* | 1.69  (5, 45) | 0.16 |
| Condition/Dose × Strain × Sex | 0.34  (5, 90) | 0.89 |  |  |  |  | 1.82 (5,90) | 0.12 |  |  |  |  |
| **Time** | 19.38  (2, 36) | <0.001* | 3.64  (2, 18) | 0.046* | 20.90 (2, 18) | <0.001* | 16.71 (2,36) | <0.001* | 12.81  (2, 18) | <0.001* | 6.57  (2, 18) | 0.01* |
| Time × Strain | 3.30  (2, 36) | 0.048* |  |  |  |  | 3.78 (2,36) | 0.03* |  |  |  |  |
| Time × Sex | 1.61  (2, 36) | 0.21 | 0.08  (2, 18) | 0.92 | 3.94  (2, 18) | 0.04* | 1.25 (2,36) | 0.30 | 1.44  (2, 18) | 0.26 | 0.41  (2, 18) | 0.67 |
| Time × Strain × Sex | 2.00  (2, 36) | 0.15 |  |  |  |  | 0.78 (2,36) | 0.47 |  |  |  |  |
| Condition/Dose × Time | 6.44  (10, 180) | <0.001* | 1.80  (10, 90) | 0.07 | 6.24  (10, 90) | <0.001* | 3.54 (10,180) | <0.001* | 3.59  (10, 90) | <0.001* | 1.13  (10, 90) | 0.35 |
| Condition/Dose × Time × Strain | 1.77  (10, 180) | 0.07 |  |  |  |  | 0.71 (10,180) | 0.71 |  |  |  |  |
| Condition/Dose × Time × Sex | 0.82  (10, 180) | 0.61 | 0.31  (10, 90) | 0.98 | 1.11  (10, 90) | 0.36 | 2.08 (10,180) | 0.03* | 2.29  (10, 90) | 0.02* | 1.30  (10, 90) | 0.24 |
| Condition/Dose × Time × Strain × Sex | 0.64  (10, 180) | 0.78 |  |  |  |  | 1.31 (10,180) | 0.23 |  |  |  |  |

Supplementary Table 1. Repeated ANOVA results for effects of strain (Sprague Dawley vs. Wistar), sex (males vs. females), time (time behavioral measures were determined), and Condition/Dose [(a) Vapor Condition 1-5 or (b) IP dose 1-20 mg/kg or vehicle conditions] on TF latency (%MPE). *Asterisks indicate a p-value below alpha significance level set at 0.05.

|  | Temperature (°C) | | | | | | | | | | | |
| --- | --- | --- | --- | --- | --- | --- | --- | --- | --- | --- | --- | --- |
|  | (a) Vapor Exposure | | | | | | (b) IP injection | | | | | |
|  | all | | SD | | Wistar | | all | | SD | | Wistar | |
|  | F (df) | p | F (df) | p | F (df) | p | F (df) | p | F (df) | p | F (df) | p |
| **Strain** | 4.00  (1, 18) | 0.06 |  |  |  |  | 2.23  (1, 18) | 0.15 |  |  |  |  |
| **Sex** | 23.89  (1, 18) | <0.001* | 19.31  (1, 9) | 0.002* | 9.52  (1, 9) | 0.01* | 0.28  (1, 18) | 0.60 | 0.02  (1, 9) | 0.88 | 1.37  (1, 9) | 0.27 |
| Strain × Sex | 0.19  (1, 18) | 0.67 |  |  |  |  | 0.62  (1, 18) | 0.44 |  |  |  |  |
| **Condition/Dose** | 115.93  (5, 90) | <0.001* | 78.90  (5, 45) | <0.001* | 50.46  (5, 45) | <0.001* | 129.86  (5, 90) | <0.001* | 89.28  (5, 45) | <0.001* | 51.56  (5, 45) | <0.001* |
| Condition/Dose × Strain | 2.22  (5, 90) | 0.06 |  |  |  |  | 1.70  (5, 90) | 0.14 |  |  |  |  |
| Condition/Dose × Sex | 1.70  (5, 90) | 0.14 | 0.89  (5, 45) | 0.50 | 1.54  (5, 45) | 0.20 | 3.54  (5, 90) | 0.01 | 2.78  (5, 45) | 0.03 | 2.47  (5, 45) | 0.046* |
| Condition/Dose × Strain × Sex | 1.00  (5, 90) | 0.43 |  |  |  |  | 1.63  (5, 90) | 0.16 |  |  |  |  |
| **Time** | 17.44  (4, 72) | <0.001* | 17.76  (4, 36) | <0.001* | 6.01  (4, 36) | <0.001* | 150.32  (4, 72) | <0.001* | 79.35  (4, 36) | <0.001* | 72.18  (4, 36) | <0.001* |
| Time × Strain | 1.75  (4, 72) | 0.15 |  |  |  |  | 1.36  (4, 72) | 0.26 |  |  |  |  |
| Time × Sex | 5.36  (4, 72) | <0.001* | 2.67  (4, 36) | 0.048* | 5.70  (4, 36) | 0.001* | 3.38  (4, 72) | 0.01 | 1.28  (4, 36) | 0.30 | 2.41  (4, 36) | 0.07 |
| Time × Strain × Sex | 4.20  (4, 72) | 0.004* |  |  |  |  | 0.29  (4, 72) | 0.88 |  |  |  |  |
| Condition/Dose × Time | 11.73  (20, 360) | <0.001* | 6.30  (20, 180) | <0.001* | 8.05  (20, 180) | <0.001* | 13.91  (20, 360) | <0.001* | 7.89  (20, 180) | <0.001* | 9.11  (20, 180) | <0.001* |
| Condition/Dose × Time × Strain | 2.76  (20, 360) | <0.001* |  |  |  |  | 2.86  (20, 360) | <0.001* |  |  |  |  |
| Condition/Dose × Time × Sex | 1.66  (20, 360) | 0.04* | 0.79  (20, 180) | 0.72 | 1.81  (20, 180) | 0.02* | 1.44  (20, 360) | 0.10 | 0.64  (20, 180) | 0.88 | 2.39  (20, 180) | 0.001* |
| Condition/Dose × Time × Strain × Sex | 1.02  (20, 360) | 0.44 |  |  |  |  | 1.26  (20, 360) | 0.20 |  |  |  |  |

Supplementary Table 2. Repeated ANOVA results for effects of strain (Sprague Dawley vs. Wistar), sex (males vs. females), time (time behavioral measures were determined), and Condition/Dose [(a) Vapor Condition 1-5 or (b) IP dose 1-20 mg/kg or vehicle conditions] on temperature. *Asterisks indicate a p-value below alpha significance level set at 0.05.

|  | Break points | | | | | | | | | | | |
| --- | --- | --- | --- | --- | --- | --- | --- | --- | --- | --- | --- | --- |
|  | (a) Vapor Exposure | | | | | | (b) IP injection | | | | | |
|  | all | | SD | | Wistar | | all | | SD | | Wistar | |
|  | F (df) | p | F (df) | p | F (df) | p | F (df) | p | F (df) | p | F (df) | p |
| **Strain** | 8.96  (1, 18) | 0.01 |  |  |  |  | 16.34  (1, 18) | <0.001* |  |  |  |  |
| **Sex** | 1.65  (1, 18) | 0.21 | 4.98  (1, 9) | 0.05 | 0.27  (1, 9) | 0.61 | 0.35  (1, 18) | 0.56 | 0.02  (1, 9) | 0.88 | 1.49  (1, 9) | 0.25 |
| Strain × Sex | 3.98  (1, 18) | 0.06 |  |  |  |  | 0.09  (1, 18) | 0.76 |  |  |  |  |
| **Condition/Dose** | 38.08  (5, 90) | <0.001* | 20.20  (5, 45) | <0.001* | 20.57  (5, 45) | <0.001* | 17.69  (5, 90) | <0.001* | 8.39  (5, 45) | <0.001* | 13.20  (5, 45) | <0.001* |
| Condition/Dose × Strain | 2.53  (5, 90) | 0.03 |  |  |  |  | 1.21  (5, 90) | 0.31 |  |  |  |  |
| Condition/Dose × Sex | 1.13  (5, 90) | 0.35 | 1.02  (5, 45) | 0.41 | 1.05  (5, 45) | 0.40 | 1.52  (5, 90) | 0.19 | 1.56  (5, 45) | 0.19 | 1.16  (5, 45) | 0.35 |
| Condition/Dose × Strain × Sex | 0.94  (5, 90) | 0.46 |  |  |  |  | 1.42  (5, 90) | 0.22 |  |  |  |  |

Supplementary Table 3. Repeated ANOVA results for effects of strain (Sprague Dawley vs. Wistar), sex (males vs. females), and Condition/Dose [(a) Vapor Condition 1-5 or (b) IP dose 1-20 mg/kg or vehicle conditions] on PR break points. *Asterisks indicate a p-value below alpha significance level set at 0.05.

|  | Response Rate | | | | | | | | | | | |
| --- | --- | --- | --- | --- | --- | --- | --- | --- | --- | --- | --- | --- |
|  | (a) Vapor Exposure | | | | | | (b) IP injection | | | | | |
|  | all | | SD | | Wistar | | all | | SD | | Wistar | |
|  | F (df) | p | F (df) | p | F (df) | p | F (df) | p | F (df) | p | F (df) | p |
| **Strain** | 1.09  (1, 18) | 0.31 |  |  |  |  | 3.63  (1, 18) | 0.07 |  |  |  |  |
| **Sex** | 2.38  (1, 18) | 0.14 | 0.55  (1, 9) | 0.48 | 2.20  (1, 9) | 0.17 | 4.81  (1, 18) | 0.04* | 6.63  (1, 9) | 0.03* | 0.29  (1, 9) | 0.60 |
| Strain × Sex | 0.18  (1, 18) | 0.67 |  |  |  |  | 2.03  (1, 18) | 0.17 |  |  |  |  |
| **Condition/Dose** | 28.87  (5, 90) | <0.001* | 22.67  (5, 45) | <0.001* | 9.79  (5, 45) | <0.001* | 7.40  (5, 90) | <0.001* | 5.86  (5, 45) | <0.001* | 2.75  (5, 45) | 0.03* |
| Condition/Dose × Strain | 0.70  (5, 90) | 0.62 |  |  |  |  | 0.46  (5, 90) | 0.80 |  |  |  |  |
| Condition/Dose × Sex | 1.07  (5, 90) | 0.38 | 0.36  (5, 45) | 0.87 | 0.90  (5, 45) | 0.49 | 0.94  (5, 90) | 0.46 | 0.50  (5, 45) | 0.77 | 1.06  (5, 45) | 0.39 |
| Condition/Dose × Strain × Sex | 0.32  (5, 90) | 0.90 |  |  |  |  | 0.76  (5, 90) | 0.58 |  |  |  |  |

Supplementary Table 4. Repeated ANOVA results for effects of strain (Sprague Dawley vs. Wistar), sex (males vs. females), and Condition/Dose [(a) Vapor Condition 1-5 or (b) IP dose 1-20 mg/kg or vehicle conditions] on PR response rates. *Asterisks indicate a p-value below alpha significance level set at 0.05.

|  | Plasma THC | | | | | | | | | | | |
| --- | --- | --- | --- | --- | --- | --- | --- | --- | --- | --- | --- | --- |
|  | (a) Vapor Exposure | | | | | | (b) IP injection | | | | | |
|  | all | | SD | | Wistar | | all | | SD | | Wistar | |
|  | F (df) | p | T (df) | p | T (df) | p | F (df) | p | T (df) | p | T (df) | p |
| **Strain** | 3.65  (1, 17) | 0.07 |  |  |  |  | 0.0004  (1, 17) | 0.99 |  |  |  |  |
| **Sex** | 3.37  (1, 17) | 0.08 | 1.59  (8) | 0.15 | 1.15  (9) | 0.28 | 5.337  (1, 17) | 0.03* | 2.43  (9) | 0.04* | 1.04  (8) | 0.33 |
| Strain × Sex | 0.002  (1, 17) | 0.96 |  |  |  |  | 0.38  (1, 17) | 0.55 |  |  |  |  |

Supplementary Table 5. Two-way ANOVA results for effects of strain (Sprague Dawley vs. Wistar) and sex (males vs. females) on plasma THC, and post-hoc unpaired t-test results in individual strains. *Asterisks indicate a p-value below alpha significance level set at 0.05.

|  |  |  | Temperature (∆*°*C) | | Tail Flick (%MPE) | |
| --- | --- | --- | --- | --- | --- | --- |
| **Vapor Condition** | **Strain** | **Sex** | *Peak Effect* | *Time to peak effect* | *Peak Effect* | *Time to peak effect* |
| 1 | Wistar | **all** | **-0.05 ± 0.0** | **150 (36.4)** | **15.9 ± 7.8** | **60 (36.4)** |
|  |  | Male | -0.02 ± 0.1 | 150 (66.7) | 11.3 ± 10.2 | 300 (33.3) |
|  |  | Female | -0.09 ± 0.1 | 30 (60.0) | 21.5 ± 12.8 | 120 (40.0) |
|  | Sprague-Dawley | **all** | **-0.05 ± 0.0*** | **30 (45.5)** | **18.1 ± 4.9** | **60 (45.5)** |
|  |  | Male | -0.06 ± 0.0 | 300 (50.0) | 14.7 ± 6.7 | 300 (50.0) |
|  |  | Female | -0.03 ± 0.1 | 60 (60.0) | 22.2 ± 7.4 | 60 (60.0) |
| 2 | Wistar | **all** | **-0.21 ± 0.1*** | **150 (45.5)** | **10.2 ± 6.7** | **60 (54.5)** |
|  |  | Male | -0.19 ± 0.1 | 150 (50.0) | 10.4 ± 6.9 | 60 (50.0) |
|  |  | Female | -0.25 ± 0.1 | 150 (40.0) | 9.8 ± 13.4 | 60 (60.0) |
|  | Sprague-Dawley | **all** | **-0.16 ± 0.0*** | **30 (63.6)** | **26.5 ± 10.2** | **60 (45.5)** |
|  |  | Male | -0.16 ± 0.0* | 30 (66.7) | 38.4 ± 17.2 | 60 (50.0) |
|  |  | Female | -0.17 ± 0.1 | 30 (60.0) | 12.3 ± 6.0 | 120 (40.0) |
| 3 | Wistar | **all** | **-0.31 ± 0.0*** | **300 (45.5)** | **5.7 ± 4.9** | **60 (54.5)** |
|  |  | Male | -0.38 ± 0.1* | 300 (66.7) | 3.2 ± 7.1 | 60 (50.0) |
|  |  | Female | -0.23 ± 0.1* | 30 (40.0) | 8.7 ± 7.3 | 60 (60.0) |
|  | Sprague-Dawley | **all** | **-0.23 ± 0.0*** | **30 (36.4)** | **12.7 ± 5.2** | **60 (72.7)** |
|  |  | Male | -0.25 ± 0.0* | 30 (50.0) | 13.9 ± 6.5 | 60 (83.3) |
|  |  | Female | -0.21 ± 0.1* | 150 (60.0) | 11.2 ± 9.2 | 60 (60.0) |
| 4 | Wistar | **all** | **-0.41 ± 0.0*** | **60 (54.5)** | **58.4 ± 12.4*** | **60 (72.7)** |
|  |  | Male | -0.31 ± 0.0* | 60 (66.7) | 44.1 ± 18.6 | 60 (66.7) |
|  |  | Female | -0.54 ± 0.0* | 60 (40.0) | 75.6 ± 13.7* | 60 (80.0) |
|  | Sprague-Dawley | **all** | **-0.46 ± 0.1*** | **60 (45.5)** | **37.8 ± 8.0*** | **60 (81.8)** |
|  |  | Male | -0.48 ± 0.1* | 30 (50.0) | 25.1 ± 7.0 | 60 (100.0) |
|  |  | Female | -0.44 ± 0.0* | 60 (60.0) | 53.1 ± 13.0 | 60 (60.0) |
| 5 | Wistar | **all** | **-0.70 ± 0.1*** | **60 (63.6)** | **74.7 ± 9.5*** | **60 (81.8**) |
|  |  | Male | -0.72 ± 0.1* | 60 (50.0) | 53.7 ± 11.7* | 60 (83.3) |
|  |  | Female | -0.67 ± 0.1* | 60 (80.0) | 99.8 ± 0.2* | 60 (80.0) |
|  | Sprague-Dawley | **all** | **-0.54 ± 0.0*** | **60 (54.5)** | **66.6 ± 7.7*** | **120 (54.5)** |
|  |  | Male | -0.51 ± 0.0* | 60 (50.0) | 63.4 ± 10.2* | 120 (50.0) |
|  |  | Female | -0.58 ± 0.1* | 60 (60.0) | 70.5 ± 12.9* | 120 (60.0) |

Supplementary Table 6. Time course of peak effects of THC vapor exposure. Temperature changes (∆) from baseline. Data for maximum ∆ temperature and latency are denoted as Mean ± SEM. Time to peak effect are denoted as Mode (% subjects reaching peak at that time). Temperature was measured between 0-300 minutes; tail flick was measured between 60-300 min.* asterisk denotes a significant difference from Vehicle (Dunnett’s).

|  |  |  | Temperature (∆*°*C) | | | Tail Flick (%MPE) | |
| --- | --- | --- | --- | --- | --- | --- | --- |
| **Dose (IP)** | **Strain** | **Sex** | *Peak Effect* | *Time to peak effect* | *Peak Effect* | | *Time to peak effect* |
| 1 mg/kg | Wistar | **all** | **-0.39 ± 0.0*** | **150 (63.6)** | **18.7 ± 4.8*** | | **15 (45.5)** |
|  |  | Male | -0.34 ± 0.0^$^ | 150 (83.3) | 21.9 ± 6.9 | | 15 (66.7) |
|  |  | Female | -0.44 ± 0.0 | 150 (40.0) | 14.9 ± 7.1 | | 60 (60.0) |
|  | Sprague-Dawley | **all** | **-0.41 ± 0.0*** | **150 (36.4)** | **9.7 ± 6.0** | | **15 (45.5)** |
|  |  | Male | -0.40 ± 0.0* | 150 (50.0) | 8.3 ± 7.8 | | 15 (50.0) |
|  |  | Female | -0.42 ± 0.0 | 300 (60.0) | 11.4 ± 10.2 | | 15 (40.0) |
| 3 mg/kg | Wistar | **all** | **-0.70 ± 0.1*** | **300 (81.8)** | **64.8 ± 12.6*** | | **60 (45.5)** |
|  |  | Male | -0.64 ± 0.1* | 300 (66.7) | 47.1 ± 18.9 | | 60 (50.0) |
|  |  | Female | -0.77 ± 0.1* | 300 (100.0) | 86.1 ± 11.3* | | 300 (40.0) |
|  | Sprague-Dawley | **all** | **-0.89 ± 0.1*** | **150 (63.6)** | **55.4 ± 11.2*** | | **300 (63.6)** |
|  |  | Male | -0.80 ± 0.1* | 150 (100.0) | 54.5 ± 15.1 | | 300 (66.7) |
|  |  | Female | -0.99 ± 0.1* | 300 (80.0) | 56.6 ± 18.7* | | 300 (60.0) |
| 5.6 mg/kg | Wistar | **all** | **-0.73 ± 0.1*** | **150 (81.8)** | **60.4 ± 9.9*** | | **300 (45.5)** |
|  |  | Male | -0.57 ± 0.0* | 150 (100.0) | 46.7 ± 13.5 | | 300 (50.0) |
|  |  | Female | -0.92 ± 0.1* | 150 (60.0) | 76.8 ± 11.8* | | 60 (60.0) |
|  | Sprague-Dawley | **all** | **-0.96 ± 0.1*** | **300 (63.6)** | **77.4 ± 10.2*** | | **60 (54.5)** |
|  |  | Male | -0.89 ± 0.1* | 300 (66.7) | 65.4 ± 16.5* | | 300 (83.3) |
|  |  | Female | -1.06 ± 0.1* | 300 (60.0) | 91.7 ± 8.3* | | 60 (100.0) |
| 10 mg/kg | Wistar | **all** | **-0.98 ± 0.1*** | **300 (45.5)** | **77.3 ± 8.9*** | | **300 (36.4)** |
|  |  | Male | -0.97 ± 0.1* | 300 (50.0) | 80.4 ± 8.9* | | 60 (50.0) |
|  |  | Female | -0.98 ± 0.2* | 90 (40.0) | 73.6 ± 17.5 | | 300 (40.0) |
|  | Sprague-Dawley | **all** | **-0.97 ± 0.1*** | **300 (63.6)** | **63.3 ± 9.9*** | | **60 (54.5)** |
|  |  | Male | -0.91 ± 0.1* | 300 (66.7) | 44.9 ± 12.5 | | 60 (50.0) |
|  |  | Female | -1.05 ± 0.1* | 300 (60.0) | 85.5 ± 8.9* | | 60 (60.0) |
| 20 mg/kg | Wistar | **all** | **-0.90 ± 0.1*** | **150 (63.6)** | **85.9 ± 8.6*** | | **15 (45.5)** |
|  |  | Male | -0.86 ± 0.1* | 150 (83.3) | 87.9 ± 9.5* | | 60 (50.0) |
|  |  | Female | -0.95 ± 0.1* | 150 (40.0) | 83.6 ± 16.4* | | 15 (60.0) |
|  | Sprague-Dawley | **all** | **-0.99 ± 0.1*** | **300 (54.5)** | **87.4 ± 8.2*** | | **60 (45.5)** |
|  |  | Male | -0.91 ± 0.1* | 300 (50.0) | 77.7 ± 14.2* | | 60 (33.3) |
|  |  | Female | -1.09 ± 0.1* | 300 (60.0) | 99.2 ± 0.8* | | 60 (60.0) |

Supplementary Table 7. Time course of peak effects of IP THC. Temperature changes ∆ from baseline. Data for maximum ∆ temperature and latency are denoted as Mean ± SEM. Time to peak effect are denoted as Mode (Percent subjects with that peak time). * asterisk denotes a significant difference from Vehicle.
